# A conserved ribosomal protein has entirely dissimilar structures in different organisms

**DOI:** 10.1101/2022.09.21.508910

**Authors:** Léon Schierholz, Charlotte R. Brown, Karla Helena-Bueno, Vladimir N. Uversky, Robert P. Hirt, Jonas Barandun, Sergey V. Melnikov

## Abstract

Ribosomes from different species can markedly differ in their composition by including dozens of ribosomal proteins that are unique to specific lineages but absent in others. However, it remains unknown how ribosomes acquire and specialize new proteins throughout evolution. Here, to help answer this question, we describe the evolution of the ribosomal protein msL1/msL2 that was recently found in ribosomes from the parasitic microorganism clade, microsporidia. We first show that this protein has a conserved location in the ribosome but entirely dissimilar structures in different organisms: in each of the analyzed species, msL1/msL2 exhibits an altered secondary structure, an inverted orientation of the N- and C-termini on the ribosomal binding surface, and a completely transformed three-dimensional fold. We then show that this evolutionary fold switching is likely caused by changes in the msL1/msL2-binding site in the ribosome; specifically, by variations in microsporidian rRNA. These observations allow us to infer an evolutionary scenario in which a small, positively-charged, *de novo*-born unfolded protein was first captured by rRNA to become part of the ribosome and subsequently underwent complete fold switching to optimize its binding to its evolving ribosomal binding site. Overall, our work provides a striking example of how a protein can switch its fold in the context of a complex biological assembly while retaining its specificity for its molecular partner. This finding will help us better understand the origin and evolution of new protein components of complex molecular assemblies – thereby enhancing our ability to engineer biological molecules, identify protein homologs, and peer into the history of life on Earth.

## INTRODUCTION

Because ribosomes are present in every organism and are thought to have originated over 3.9 billion years ago, at the very dawn of life, their structures are widely used to gain insights into past events that are irreparably lost to our direct observations(Bowman, et al. 2020). Some of these events include the origin of species(Woese and Fox 1977; Schmidt and Relman 1994; Fox, et al. 2012; Petrov, et al. 2015), the rise of catalytic RNAs(Fox, et al. 2012; Bose, et al. 2022), the origin of the genetic code and protein chirality(Fox 2010; Polikanov, et al. 2015; Melnikov, et al. 2019), as well as adaptations of biological molecules to cellular compartmentalization(Melnikov, et al. 2015; Melnikov, et al. 2020) or new environments(Di Giulio 2003; Khachane, et al. 2005; Wang, et al. 2006; Kimura, et al. 2007; Kimura, et al. 2013; Hu and Lercher 2021; Hu, et al. 2022). Studying these evolutionary processes is important beyond genuine curiosity aiming to understand our roots; this knowledge is also essential for synthetic biology to help improve natural enzymes or engineer new life from scratch.

Many principles of the molecular evolution of ribosomes have been revealed through the determination of their atomic structures. These structures revealed, for example, that the ribosome’s enzymatic core consists of an RNA dimer, hinting at a dimeric primordial RNA as the ribosome ancestor(Belousoff, et al. 2010). Later, the discovery of RNA folding mechanisms allowed to infer the order of events that transformed a relatively small primordial rRNA into one of the most complex molecular assemblies observed in living cells today (Bokov and Steinberg 2009; Petrov, et al. 2014).

While structural studies of ribosomes have made it possible to infer the origin, evolution, and functional specialization of rRNA across species, we know little about the origin of ribosomal proteins. If we compare, for example, ribosomes in *Escherichia coli* and humans, we find that they share a set of 33 conserved proteins. However, in addition, these ribosomes contain an array of seemingly unrelated proteins, including 21 bacteria-specific proteins in *E. coli* and 47 archaeo-eukaryotic proteins in humans(Melnikov, et al. 2012), with many of these proteins preforming essential roles in protein synthesis.

Despite this strikingly dissimilar protein content, the process by which ribosomes acquire and adapt new proteins in different species remains unknown(Kovacs, et al. 2017; Alvarez-Carreno, et al. 2021; Alvarez-Carreno, et al. 2022). Did lineage-specific ribosomal proteins first emerge as primitive peptides that gradually transformed into globular proteins? Or did ribosomes capture existing globular proteins through a gain-of-function event? How do these proteins change and coevolve with ribosomes throughout evolutionary time? While these are huge and complicated questions that cannot be comprehensively addressed in any single study, analysis of ribosome structures and sequences from different organisms can shine light on possible evolutionary scenarios.

Recently, an unexpected opportunity to explore these questions arose from structural studies of ribosomes from the group of fungal parasites known as microsporidia (Barandun, et al. 2019; Ehrenbolger, et al. 2020; Nicholson, et al. 2022). These studies revealed that microsporidian ribosomes possess a unique ribosomal protein, msL1/msL2, which appears to compensate for the rRNA size reduction in microsporidian species. Compared to other studied eukaryotes, most microsporidian species possess severely truncated rRNA which is devoid of the characteristic rRNA expansion segments that distinguish eukaryotes from bacteria. Also, microsporidians have truncations in most ribosomal proteins (Barandun, et al. 2019; Ehrenbolger, et al. 2020; Nicholson, et al. 2022). Proteins msL1/msL2 appear to compensate for some of these truncations, including the absence of the N-terminus of protein uL23 and the loss of rRNA expansion segments ES19 and ES31 in the 25S rRNA: msL1/msL2 fill the voids in the truncated ribosome structure, mimicking the N-terminus of uL23 and also maintaining contacts between rRNA and ribosomal proteins that are supported by ES19 and ES31 in other eukaryotic ribosomes. Therefore, the presence of msL1/msL2 likely allows microsporidia to maintain the normal ribosome biogenesis despite truncations in ribosomal proteins and rRNA.

First discovered in the parasite *Vairimorpha necatrix*, this protein was annotated as msL1-*Vn*; its homologs were subsequently found in 10 microsporidian species, including the parasite *Encephalitozoon cuniculi* (Barandun, et al. 2019; Nicholson, et al. 2022). Later, however, an investigation of the structure of *E. cuniculi* ribosomes revealed that the msL1-*Vn* binding site was occupied by a structurally dissimilar protein, which was annotated as msL2-*Ec* (Nicholson, et al. 2022). The study of *E. cuniculi* ribosomes, however, has overlooked that the proteins msL1-*Vn* and msL2-*Ec* were previously described as distant homologs following an iterative search of msL1-*Vn* homologs (Barandun, et al. 2019). This homology was easy to overlook because msL1-*Vn* and msL2-*Ec* have highly dissimilar structures, with a root-mean-square deviation (RMSD) of approximately 11.4 Å between their backbone residues, compared with a typical RMSD of less than 2.4 Å for close homologs and approximately 4.5 Å for highly distant homologs.

Here, we describe this structural anomaly, in which a seemingly conserved ribosomal protein has completely transformed its three-dimensional structure in one species relative to another. We provide evolutionary insight into the mechanism of this structural shift, likely driven by its adaptation to rRNA reduction in microsporidian parasites. Furthermore, we show that while msL1-*Vn* and msL2-*Ec* share 41% sequence similarity, their conserved residues form dissimilar contacts with the *E. cuniculi* and *V. necatrix* ribosomes. Thus, despite being classified as homologs based on canonical sequence analysis and binding to the same ribosomal helix H52, msL1-*Vn* and msL2-*Ec* have entirely dissimilar three-dimensional structures, binding orientations, and binding modes to the ribosome.

## RESULTS

### The msL1/msL2 fold switch in microsporidian species

To better understand how the proteins msL1-*Vn* and msL2-*Ec* could have changed their fold during their evolution, we first compared their structures and ribosome attachment mechanisms. Both msL1-*Vn* and msL2-*Ec* were clearly visualized in the cryo-EM structures of their respective ribosomes (PDB accession codes 6RM3 and 7QEP) and display unique and distinct folds compared to each other and to AlphaFold predictions(Barandun, et al. 2019; Nicholson, et al. 2022) (**Fig. 1a**). Although the two proteins are of identical length (72 amino acids), share 41% sequence similarity, and some secondary structure similarity, namely two short α-helical segments (residues 23-40 and 58-68) the overall folds are very dissimilar with a backbone RMSD value of 11.4 Å.

**Figure 1.**
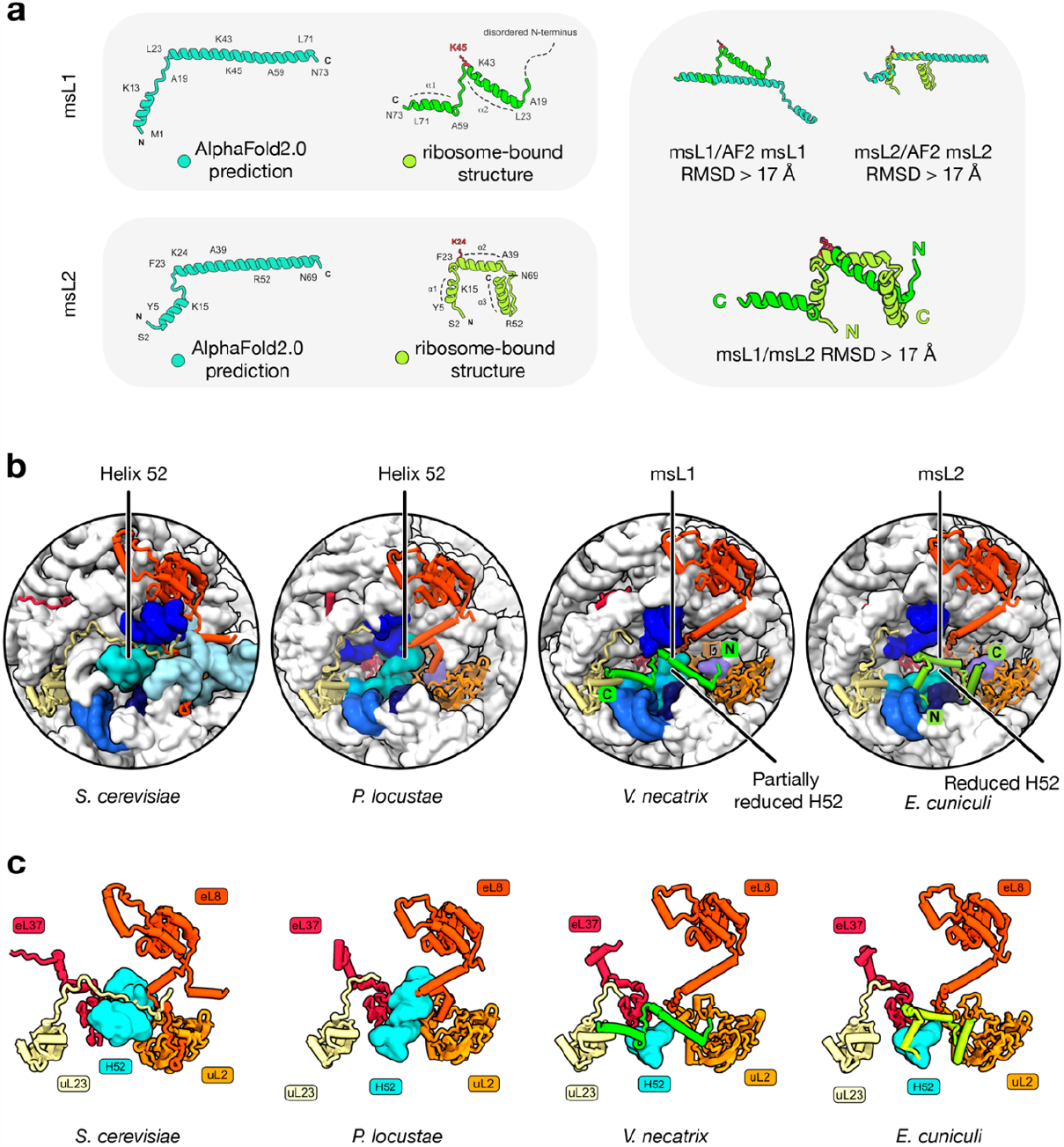
The msL1/msL2 fold switching in microsporidian ribosomes. **a)** The left panels compare structures of two homologous ribosomal proteins, msL1-*Vn* and msL2-*Ec*, as predicted by AlphaFold2 (labelled as “AlphaFold2.0 prediction”) or observed in the structures of *V. necatrix* and *E. cuniculi* ribosomes (labelled as “ribosome-bound structure”), respectively. The right panels show superpositions between the AlphaFold2-predicted and experimentally determined structures of msL1 and msL2 as well as the superposition of the experimentally determined structures of msL1 and msL2 between each other, providing the RMSD values for each of these superpositions. **b)** Zoom-in views of ribosome structures compare the binding sites for msL1*-Vc* and msL2*-Ec* and corresponding ribosomal sites in *S. cerevisiae* and *P. locustae*. In microsporidian ribosomes compared to ribosomes from other eukaryotes, the helix H52 is truncated. In *V. necatrix*, the void of the truncated H52 is filled with msL1-*Vn*, and in *E. cuniculi* ribosomes this void is filled with msL1-*Ec*. **c)** Side-by-side comparison of the molecular surrounding of msL1*-Vn*, msL2*-Ec* and H52 in the ribosome from different species.

We next compared the ribosomal binding mode and location of msL1-*Vn* and msL2-*Ec*. While msL1-*Vn* and msL2-*Ec* are attached to the same, although variable in size, 25S rRNA helix H52 in the large ribosomal subunit, our analysis showed that in *V. necatrix* ribosomes, the N-terminus of msL1-*Vn* is directed toward the solvent-exposed side of the ribosome, whereas in *E. cuniculi* ribosomes, the N-terminus of msL2-*Ec* is oriented in the opposite direction, toward the ribosomal core. Thus, the proteins msL1-*Vn* and msL2-*Ec* are inverted relative to each other in the ribosome structure, pointing their N- and C-termini in opposite directions (**Fig. 1b**).

Next, we aimed to understand whether the conserved residues of these proteins make conserved contacts with the ribosome. Given the extent of sequence similarity between msL1-*Vn* and msL2-*Ec* (41% similarity), we compared the molecular contacts of 21 identical and 8 similar residues with the ribosome. We found that most of these conserved residues mediated msL1-*Vn*/msL2-*Ec* attachment to the ribosome; strikingly, however, none of them bound the ribosome in a similar manner in both proteins. For example, in *V. necatrix* ribosomes, the conserved motif ^43^KIKxLKxKK^51^ of msL1-*Vn* bound the bundle of rRNA helices H5/H8/H10/H52/H53, whereas in *E. cuniculi* ribosomes the same motif was moved approximately 22 Å away, binding a different bundle, H55/H56/H66 (**Fig. 1b,c**).

Overall, our analysis revealed a vivid contrast between a relatively high conservation of msL1-*Vn*/msL2-*Ec* sequences and complete change of fold. Thus, msL1-*Vn* and msL2-*Ec* represent a new example of proteins classified as sequence homologs that adopt dissimilar folds and completely repurpose their conserved residues in the ribosome structure to retain their biological function.

### Each microsporidian lineage bears a dissimilar ortholog of msl1/msL2

To better understand the evolution of msL1/msL2 folds, we next compared msL1-*Vn* and msL2-*Ec* with their structurally uncharacterized homologs from other microsporidian species. In doing so, we sought to understand whether these uncharacterized homologs resemble msL1-*Vn* or msL2-*Ec* in ribosome attachment and possibly in fold. To address this, we first searched for msL1/msL2 homologs in the NCBI genome database using a sensitive homolog search based on hidden Markov models (**Materials and Methods, Supplementary Data S1**). This search did not detect any msL1/msL2 homologs outside of the microsporidian clade but revealed detectable msL1/msL2 homologs in 16 of the 24 characterized microsporidian species, especially in the microsporidian clades that are most distant from other eukaryotic species and exhibit the highest extent of genome reduction and rRNA reduction (**Fig. 2a-c**).

**Figure 2.**
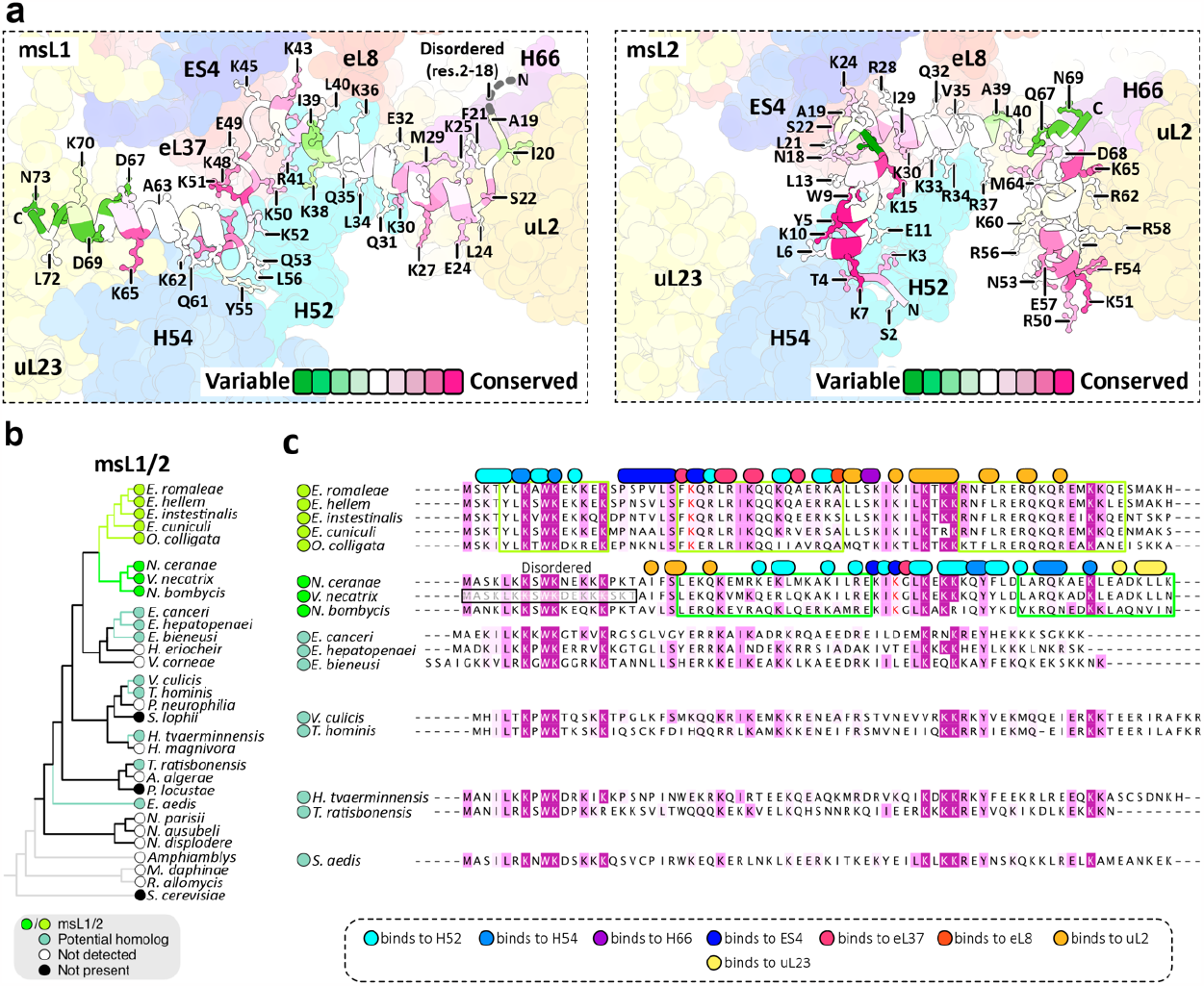
Each lineage of microsporidians encodes a dissimilar ortholog of msl1/msL2. **a)** Structures of msL1-*Vn* and msL2-*Ec* are shown in their ribosome-binding sites and colored by conservation of their residues across microsporidian species. The panel shows that the ribosome-binding residues of msL1 and msL2 include both highly conserved and highly variable residues, and that highly conserved residues of msL1/msL2 (e.g. K65) have dissimilar location in the ribosome structure in *V. necatrix* (left panel) compared to *E. cuniculi* (right panel). **b)** A ribosomal protein-based phylogenetic tree highlights microsporidian species with detectable msL1/msL2 homologs. The label “not detected” indicates that the absence msL1/msL2 homologs in corresponding microsporidian species was inferred from sequence-based homology searches. The label “not present” indicates that the molecular structure of ribosomes from the corresponding microsporidian species was determined and showed the absence of msL1/msL2 homologs in the ribosome. **c**) A multiple sequence alignment illustrates the conservation msL1/msL2 homologs from different microsporidian species. This alignment also highlights residues in msL1-*Vn* and msL2-*Ec* that are involved in ribosome binding. Each of these ribosome-binding residue is indicated as a circle or an oval above the aligned sequences and is colored by their molecular partner in the ribosome structure (e.g.). Green boxes around aligned sequences indicate helical segments of msL1-*Vn* and msL2-*Ec*.

We then assessed the sequence similarity between these homologs and msL1*-Vn* and msL2-*Ec* (**Fig. 2c**, Supplementary **Figs. 1 and 2**). We found that among *Encephalitozoon* species, msL1/msL2 homologs shared more than 95% of sequence conservation in those residues that mediate the ribosome attachment of msL2-*Ec*, suggesting that in *Encephalitozoon* species these homologs share the same ribosome attachment strategy and possibly the same fold as in msL2-*Ec*. We also found that msL1-*Vn* shared less than 48% sequence conservation in those residues that mediate ribosome attachment in msL2-*Ec*, consistent with the fact that msL1-*Vn* has a dissimilar ribosome attachment strategy and fold compared with those of msL2-*Ec*.

The most rapidly evolving residues are located in the middle of the msL1/msL2 molecules (residues 38-39 in msL1-Vn) and at the C-terminus (residues 69-73 in msL1-Vn). However, despite sequence variability, these segments have the same secondary structure (**Fig. 2a**). The most conserved residues cluster in two segments of msL1/msL2: at the N-terminus, which contains the ^7^KxKW^10^ motif that is conserved among all msL1/msL2 homologs, and in the loop in the middle of the msL1/msL2 polypeptide chain, which contains the highly conserved ^47^LKxKK^51^ motif. However, regardless of their conservation, even the most conserved residues in msL1/msL2 have entirely dissimilar surroundings in the ribosome from *E. cuniculi* compared to the ribosome from *V. necatrix* (**Fig. 2c**).

Remarkably, we found that the remaining microsporidian species carried mutations in most of the residues that are responsible for ribosome attachment of msL1-*Vn* or msL2-*Ec* (**Fig. 2a-c**). For example, the *Trachipleistophora hominis* homolog of msL1/msL2 carries mutations in approximately 66% of the ribosome-binding residues of msL1-*Vn* and 68% of the ribosome-binding residues of msL2-*Ec*. Hence, if the msL1/msL2 homolog from *T. hominis* has the same fold as msL1-*Vn* or msL2-*Ec*, it would carry mutations in most of its ribosome-binding residues. This high variability of the ribosome-binding interface suggests that each lineage of microsporidian parasites (such as *Encephalitozoon, Nosematidae, Enterospora*, and *Trachipleistophora*) may encode structurally dissimilar variants of msL1/msL2.

### The msL1/msL2 fold change is likely caused by rRNA truncation

Finally, to understand what may have caused the fold- and orientation switch between msL1-*Vn* and msL2-*Ec*, we compared the molecular surroundings of these proteins in the ribosome. We hypothesized that alterations in ribosomal structure at the binding site may have induced a coevolutionary adaptation in the msL1/msL2 protein to maintain binding. To test this idea, we first superposed the structures of *V. necatrix* and *E. cuniculi* ribosomes and found that the msL1/msL2-binding pockets are not identical, with the most prominent difference present in helix H52 of the 25S rRNA (**Fig. 3a**). This helix is three bases shorter in *E. cuniculi* compared with the *V. necatrix* ribosome. Strikingly, if the *V. necatrix* helix were the same size as in *E. cuniculi*, it would occlude 24% of the ribosome-binding surface for msL1-*Vn*, potentially explaining why msL1-*Vn*/msL2-*Ec* cannot adopt the same fold in *E. cuniculi* and *V. necatrix* ribosomes.

**Figure 3.**
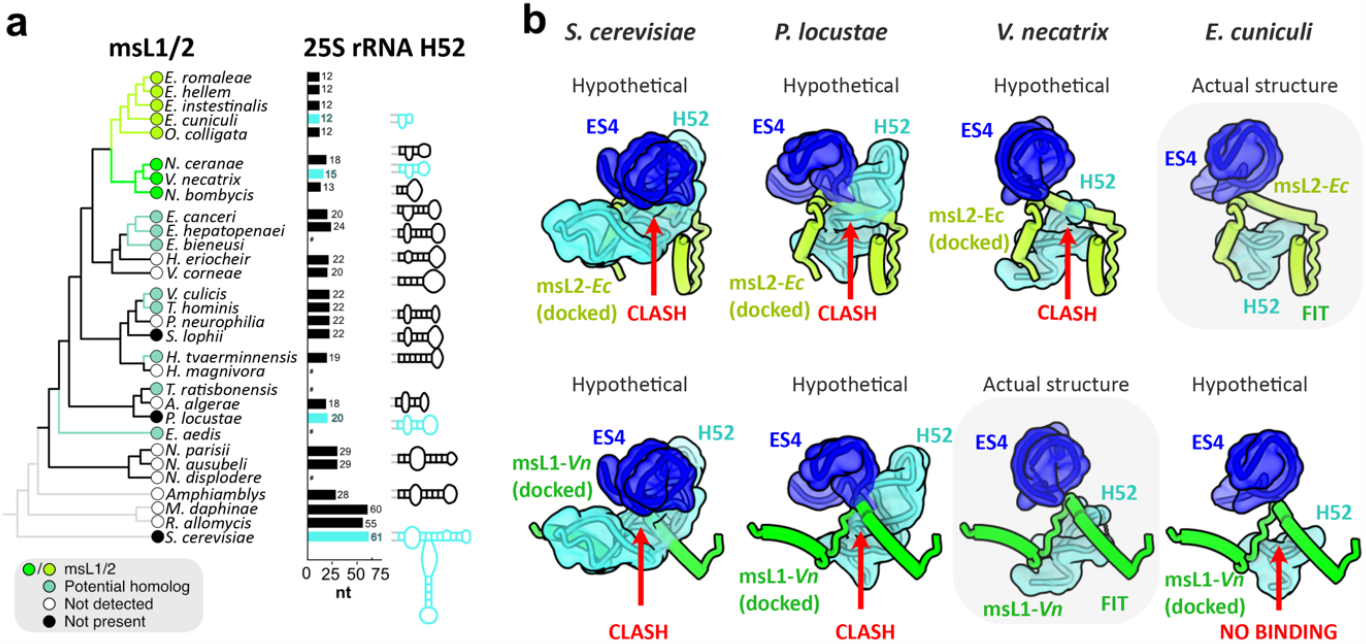
The msL1/msL2 fold change is likely caused by rRNA truncations. **a)** A ribosomal protein-based phylogenetic tree is shown with detectable msL1/msL2 homologs indicated. For species with available 5.8S-25S rRNA sequences, the length and predicted secondary structure of the helix H52 are shown next to their corresponding species. Species with missing or incomplete sequences for 5.8S-25S rRNA are highlighted by asterisks, and species with experimentally defined ribosome structures are highlighted in cyan. The label “not detected” indicates that the absence msL1/msL2 homologs in corresponding microsporidian species was inferred from sequence-based homology searches. The label “not present” indicates that the molecular structure of ribosomes from the corresponding microsporidian species was determined and showed the absence of msL1/msL2 homologs in the ribosome. **b)** The structures of msL1-*Vn* and msL2-*Ec* docked into the ribosomal environment of *S. cerevisiae, P. locustae, V. necatrix and E. cuniculi* ribosomes to illustrate potential steric clashes or loss of ribosome-binding surface in one species relative to each other. Each docking was performed by superimposing ribosome structures from *E. cuniculi* and *V. necatrix* with ribosome structures with other eukaryotic species.

Because this finding suggested that the H52 truncation was a major driver of the msL1-*Vn*/msL2-*Ec* fold switch, we next assessed the length of H52 across microsporidian species (**Fig. 3a, Supplementary Data 2**). We found that in typical eukaryotes, such as *Saccharomyces cerevisiae* and humans, H52 comprises ∼60 rRNA bases, whereas in microsporidian species H52 is reduced to 12-33 bases. We observe msL1/msL2 orthologs present in every sequenced microsporidian lineage with short H52s (12 to 18 bases), but more sporadic presence of msL1/msL2 in lineages with longer H52s (19 to 24 bases). *Paranosema locustae*, in which we did not detect an msL1/msL2 homolog, has a longer H52 (20 bases), and the previously published structure of its ribosome (PDB accession code 6ZU5) confirms the absence of msL1/msL2(Ehrenbolger, et al. 2020). Docking msL1-*Vn* and msL2-*Ec* into their ribosomal binding sites on the *P. locustae* or *S. cerevisiae* ribosome revealed that the longer H52 would not be able to accommodate msL1-*Vn* nor msL2-*Ec* (**Fig. 3b**). Collectively, our findings suggest that variations in rRNA size can indeed be a driver of the msL1-*Vn*/msL2-*Ec* fold change.

## DISCUSSION

### A conserved ribosomal protein has entirely dissimilar folds in different organisms

In this study, we capitalized on the recent advances in structural studies of microsporidian ribosomes to explore how ribosomes alter new ribosomal proteins during their evolution. This approach enabled us to identify an evolutionary fold switch in the ribosomal protein msL1/msL2 that would be impossible to observe using computational predictions or sequence analyses. We found that this ribosomal protein underwent a nearly complete transformation of its secondary and three-dimensional structure while retaining its ability to recognize the same specific ribosomal helix, H52, in one organism compared with another.

This finding has several implications for our understanding of how large macromolecular machines, including ribosomes, acquire new subunits and influence their structural fold across evolutionary time. Firstly, our work allows us to reevaluate the concept of structural mimics: structurally dissimilar proteins that occupy the same site in a given macromolecular complex from different species(Harms, et al. 2001). For example, in bacterial ribosomes, protein bL34 occupies the same site as protein eL37 in eukaryotic ribosomes(Harms, et al. 2001). Because bL34 is composed of β-sheets and eL37 of α-helices and these proteins share only 19% of sequence similarity, they are assumed to have independent evolutionary origins. However, the example of msL1/msL2 (although these proteins have twice higher sequence similarity) highlights that a lack of structural similarity does not necessarily imply an independent or even distant evolutionary origin(Harms, et al. 2001; Klein, et al. 2004; Ban, et al. 2014).

Also, our study of the msL1/msL2 protein family provides an example where the most conserved protein property may not be its sequence or structure but its position within the macromolecular assembly to which it belongs. Consequently, it is possible that for proteins acting as part of complex molecular assemblies, their position within these assemblies can serve as an additional potential indicator of their evolutionary origin.

It is important to note that, without additional experimental support, our understanding of msL1/msL2 evolution is hypothetical and does not exclude alternative evolutionary scenarios. One such scenario could involve an ancient gene duplication event of the msL1/msL2 predecessor, followed by the specialization of two paralogs and the loss of one of these isoforms in certain microsporidian species. The presence or absence of msL1 or msL2-coding genes in certain microsporidian clades could also result from horizontal gene transfer events, among other scenarios. Therefore, it will be important to explore ribosomes from other microsporidian lineages, including those from *Enterocytozoon* and *Vavraia*/*Trachipleistophora*, to better distinguish between these alternative scenarios of msL1/msL2 evolution.

### msL1/2 is an example of an evolved fold switching without loss of biological specificity

Previously, the ability of proteins to completely transform their structural fold through evolution has been studied in detail for more than a dozen proteins, and was termed as “macrotransitions”, “evolutionary metamorphosis” or “evolved fold switching” (Grishin 2001; Alva, et al. 2008; Alexander, et al. 2009; Farias-Rico, et al. 2014; Andreeva, et al. 2015; Toledo-Patino, et al. 2019; Kolodny, et al. 2021; Chakravarty, et al. 2023). One of the most striking examples of this metamorphosis was found in *Streptococcus* protein G, where the substitution of just a single amino acid, L45Y, alters eighty-five percent of protein’s secondary structure, transforming the serum-binding domain protein G_A_ into the IgG-binding domain protein G_B_(Alexander, et al. 2009). In these previously studied examples, however, the observed fold change was typically accompanied by a change in biological function of a given protein. In contrast, we showed that protein msL1/msL2 changes its fold while retaining its ability to recognize one specific partner: of all possible binding sites in a cell, this protein retains its ability to bind the specific site of the ribosome at the tip of 25S rRNA helix H52. This example shows that a protein can, in the right context, undergo evolved fold switching without function loss.

### msL1/2 is an example of fold switching likely driven by changes in its binding partner

Recent advances in protein structural prediction by AlphaFold2, ESM2, and other AI-based tools have shifted the gravity of protein studies toward Anfinsen’s folders, globular proteins whose structures are determined by their amino acid sequences(Xu and Zhang 2013; Ovchinnikov, et al. 2017; Deng, et al. 2019; Sillitoe, et al. 2019; Andreeva, et al. 2020; Gligorijevic, et al. 2021; Jumper, et al. 2021; Pereira, et al. 2021; Aderinwale, et al. 2022). Nature, however, carries a high number of protein families that either require chaperones to fold into a certain structure, or that lack globular domains altogether and remain intrinsically disordered unless they bind to other molecules(Oldfield and Dunker 2014; Uversky 2016; Pancsa, et al. 2018; Macossay-Castillo, et al. 2019). Based on recent estimates, these non-globular and intrinsically disordered proteins include more than 1,150 protein families, and more than 50% of eukaryotic proteins were shown to bear at least one long disordered segment (Tompa 2012; Wright and Dyson 2015; Contreras-Martos, et al. 2018; Kulkarni and Uversky 2018; Toto, et al. 2022). This abundance of non-globular proteins, along with their frequent omission from structural studies, raises the question: do they follow the same principle of evolutionary fold switching as their globular counterparts?

Our analysis of msL1/msL2 protein provide one possible answer to this question: msL1/msL2 is a typical non-globular protein in both *E. cuniculi* and *V. necatrix*, and our computational analysis revealed that the sequences of all known msL1/msL2 homologs are characterized by high levels of predicted intrinsic disorder (see **Supplementary Figures S1** and **S2**). The fact that the structures of msL1 and msL2 are similar when predicted by Alphafold but differ from the experimentally determined ribosome-bound structures indicates that the ribosome-bound structures of msL1/msL2 are likely defined by their contacts with the ribosome. Specifically, the rRNA helix H52 appears to promote folding of msL1/msL2 by neutralizing the positive charge of the msL1/msL2 molecule.

Overall, our analysis suggests an evolutionary scenario in which ancient microsporidians have lost one of their rRNA segments (ES31L) and evolved a new protein through *de novo* gene birth to counteract the negative impact of the rRNA truncation. As microsporidians continued to undergo degeneration of their rRNA, the new ribosomal protein underwent a fold switch to maintain its binding to the ribosome despite structural changes in how it engages with the ribosome. This scenario can be depicted by a classical Sewall Wright adaptive fitness landscape, where deleterious mutations in rRNA are compensated by the emergence and subsequent evolution of a novel ribosomal protein (**Fig. 4**). This model is consistent with previous work demonstrating that microsporidian rRNA appears to evolve faster than their ribosomal proteins(Peyretaillade, et al. 1998; Lecompte, et al. 2002; O’Mahony, et al. 2007; Melnikov, Manakongtreecheep and Soll 2018; Melnikov, Manakongtreecheep, Rivera, et al. 2018; Barandun, et al. 2019; Ehrenbolger, et al. 2020; Wadi and Reinke 2020; Jespersen, et al. 2022; Nicholson, et al. 2022). Thus, our study of msL1/msL2 provides strong evidence that, in non-globular proteins, evolutionary changes in their binding partners can serve as the major driver of an evolutionary fold switch.

**Figure 4.**
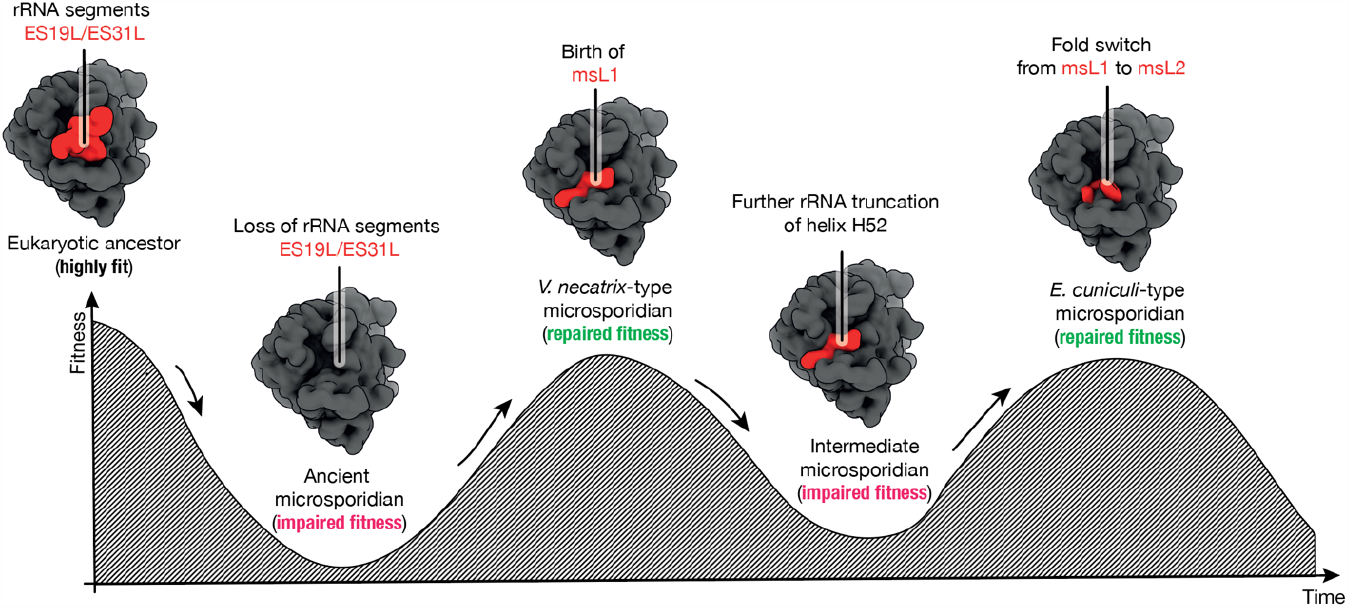
A hypothetical evolutionary scenario of the origin and fold switch of ribosomal protein msL1/msL2. The Sewall Wright’s adaptive fitness landscape shows a possible order of events in which degeneration of rRNA in microsporidian species was compensated by the origin and subsequent evolution of a new ribosomal protein. In this hypothetical order of events, microsporidian species have undergone a progressive reduction of their ribosomal RNA, which was likely caused by the accumulation of deleterious mutations in microsporidian genomes due to frequent genetic drifts. Then a new protein msL1 could originate through the spontaneous *de novo* birth of a gene that was retained in microsporidia due to its ability to counterbalance the rRNA reduction. As rRNA continued to evolve and reduce in size, protein msL1 then underwent a fold switch to yield protein msL2 with improved the affinity to its ribosomal binding site (i.e. truncated helix H52).

## MATERIALS AND METHODS

### Comparative analysis of microsporidian ribosome structures

The msL1/msL2 structures in *E. cuniculi* and *V. necatrix* ribosomes were compared using Coot (0.9.4.1) to superpose structures, measure inter- and intramolecular distances, and calculate the RMSD between protein backbone residues(Emsley, et al. 2010). The contact area between msL1/msL2 and rRNA was calculated using the CCP4 package AreaIMol 8.0.001, with the solvent sphere diameter set to 1.7Å(Hough and Wilson 2018). The figures were prepared using ChimeraX 1.2.5(Pettersen, et al. 2021). Intrinsic disorder propensities of the ribosomal proteins from the msL1/msL2 family were evaluated by the PONDR® VLXT computational tool(Romero, et al. 2001). To evaluate global disorder predisposition of a query protein, we calculated the percent of predicted disordered residues (i.e., residues with disorder scores above 0.5) and calculated the average disorder score as a protein length-normalized sum of all the per-residue disorder scores in a query protein. For further classification of these proteins, we used CH-CDF analysis(Mohan, et al. 2008; Huang, et al. 2012; Huang, et al. 2014).

### Structure prediction using AlphaFold

Structural model predictions were carried out using the AlphaFold Colab Notebook (version 2.3.2)(Mirdita 2022) using the full sequences of msL1-Vn and msL2-Ec. All modeling was performed using default parameters without any additional environmental factors, and the structures were visualized in ChimeraX 1.2.5(Pettersen, et al. 2021).

### Searching for msL1/msL2 homologs

Putative msL1/msL2 homologs were identified using two iterations of HMMER(Finn, et al. 2011), with sequences of msL1/msL2 from either *E. cuniculi* or *V. necatrix* as the initial input. For each search iteration, we used the following search options: -E 1 --domE 1 --incE 0.01 --incdomE 0.03 --seqdb uniprotrefprot. To maximize the success rate, the search was repeated using the following databases of protein sequences: Reference Proteomes, UniProtKB, SwissProt, and Ensemble. The obtained results from each of these searches were combined, reduced to remove redundant findings, and listed (**Supplementary Data 1**). The identified sequences of msL1, msL2, and their apparent homologs were then used as an input file to assess the conservation of msL1/msL2 residues using Rate4Site 2.01 with the default parameters(Pupko 2002).

### Analysis of variations in microsporidian rRNA

Due to the anomalously short length of rRNA from microsporidian species compared to other eukaryotes, many of microsporidian rRNA sequences are typically excluded from such commonly used databases of rRNA sequences such as SILVA due its “truncated sequences” filter(Quast, et al. 2013). Therefore, to analyze variations in microsporidian rRNA, microsporidian 25S rRNA sequences were consolidated using the SILVA ribosomal RNA gene database(Quast, et al. 2013), the RNAcentral database(The 2019), MicrosporidiaDB(Warrenfeltz, et al. 2018), as well as local genome databases for *V. necatrix* and *P. locustae*. These sequences were then aligned using Clustal Omega with default parameters, the resulting alignment is shown in (**Supplementary Data 2**). Where available, ribosomal structures were used to manually model the 2D structures of helix H52. These structures included the models of *S. cerevisiae* (PDB 4V88), *V. necatrix* (PDB 6RM3), *P. locustae* (PDB 6ZU5) and *E. cuniculi* (PDB 7QEP) ribosomes. For the remaining species, respective segments of the 25S rRNA alignment were used to predict 2D structures with Mfold using the RNA Folding Form feature(Zuker 2003). The sequences corresponding to the 2D structures of Helix H52 were then assessed to calculate the total length of the rRNA segment in each microsporidian species.

### Assembling a microsporidian phylogenetic tree

The microsporidian phylogeny shown in (**Fig. 2b**) and (**Fig. 3a**) was assembled based on ribosomal protein sequences. Protein sequences for *V. necatrix* and *P. locustae* were procured from local databases(Barandun, et al. 2019; Ehrenbolger, et al. 2020). Ribosomal protein sequences were retrieved by using the translated nucleotide blast (tblastn) with an E-value cutoff of 0.05 based on sequences of the *S. cerevisiae* or *V. necatrix* hits as query and the NCBI NIH and the MicrosporidiaDB(Warrenfeltz, et al. 2018) as the search databases. The protein sequences were aligned using MUSCLE(Edgar 2004), followed by trimming with trimAl(Capella-Gutierrez, et al. 2009) using the “–gappyout” option. Trimmed alignments were subsequently concatenated using FASconcat(Kuck and Meusemann 2010). The resulting phylogenetic tree was assembled with IQ-TREE and visualized using FigTree.v1.4.4 using *S. cerevisiae* as the root.

## ACKNOWLEDGEMENTS AND FUNDING SOURCES

We thank members of the Barandun and Melnikov laboratories for help with the manuscript preparation and Johan Panek (Newcastle University, UK) for his critical comments. This work was funded by the BBSRC UK Studentship BB/T008695/1 (to C.R.B.), the NUORS 2021 award (to K.H-B.), the Swedish Research Council (2019-02011), the European Research Council (ERC Starting Grant PolTube 948655), the SciLifeLab National Fellows program, and MIMS (all to J.B.) and the European Union’s Horizon 2020 research and innovation programme (the Grant Agreements No. 895166 to S.M).

