## Supplementary Information for "A conserved ribosomal protein has entirely dissimilar structures in different organisms"

*A ribosomal protein binding the same ribosomal site*

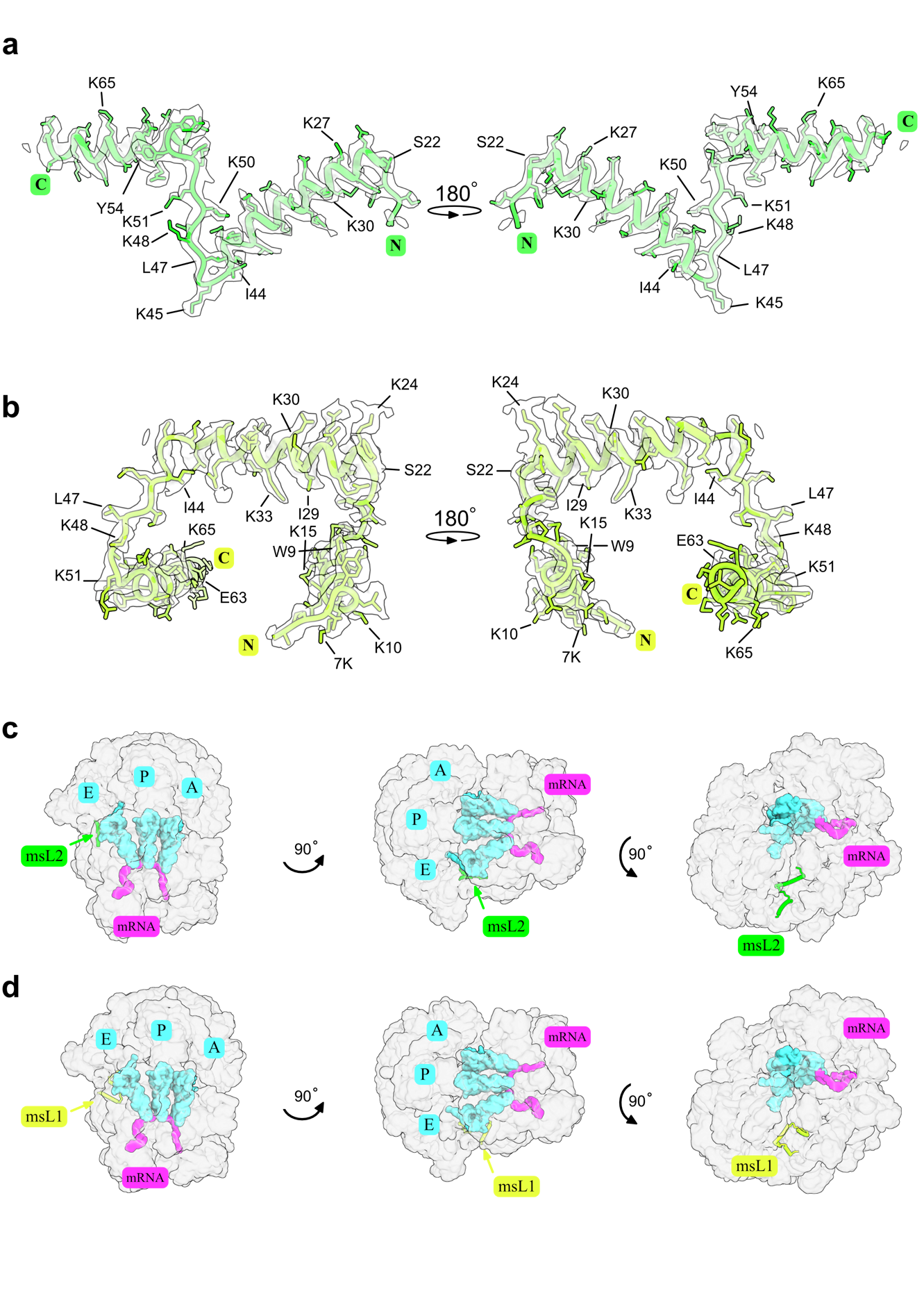

**Figure S1. The molecular structures and locations of proteins msL1 and msL2 in the structure of microsporidian ribosomes from *V. necatrix* and *E. cuniculi*, respectively.** (**a**,**b**) The cryo-EM maps and atomic models showing the structure of ribosomal protein msL1 in the ribosome from microsporidian parasites *V. necatrix* (**a**), and protein msL2 in the ribosome from microsporidian parasites E. cuniculi (**b**). (**c, d**) Structures of the ribosome shown in three orthogonal views, illustrating the location of the ribosomal protein msL2 (**c**) and msL1 (**d**) relative to the universally conserved mRNA and tRNA-binding sites of the ribosome.

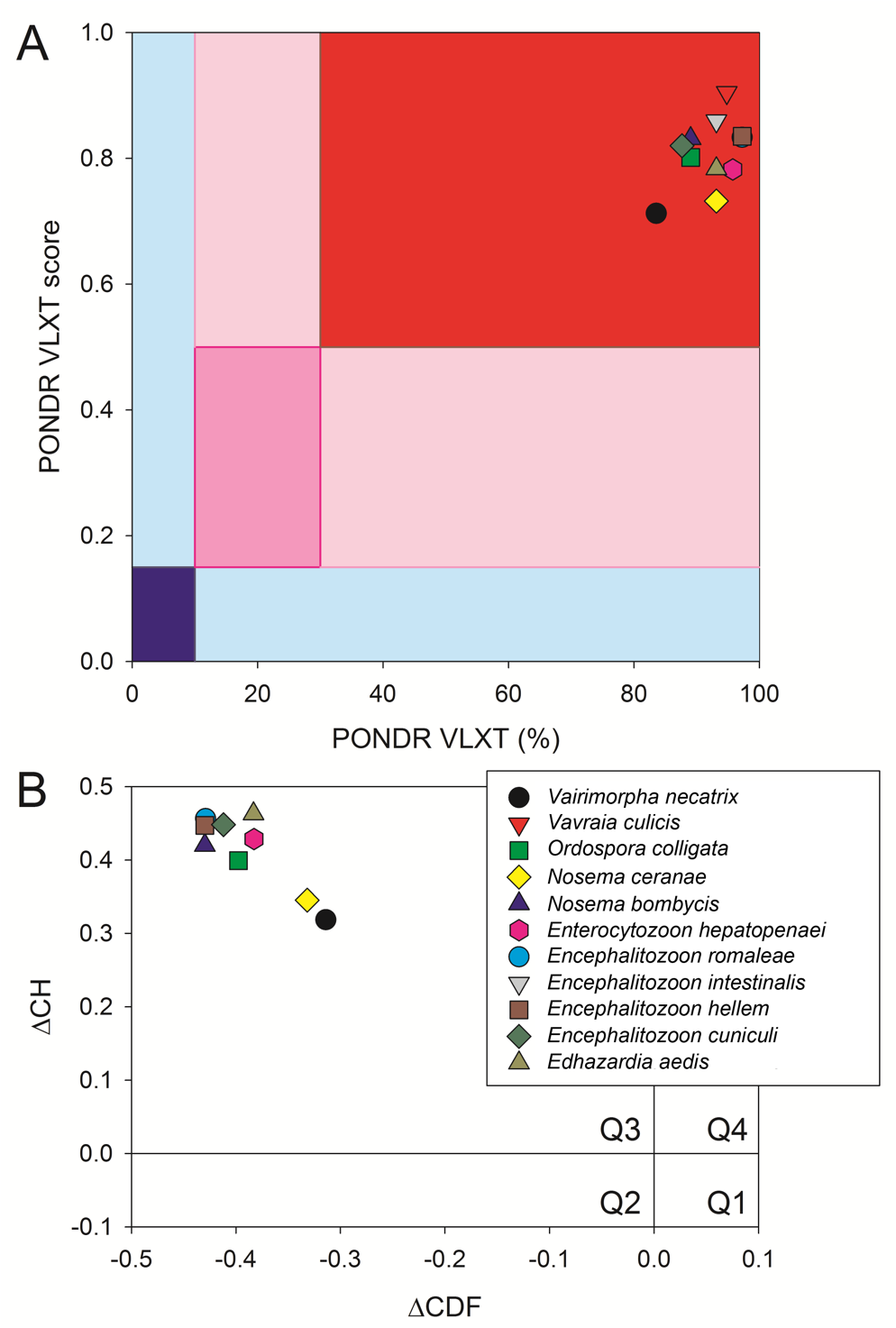

**Figure S2. Global evaluation of the intrinsic disorder predisposition of the ribosomal proteins from the msL1/msL2 family.** **A**. The PONDR® VLXT output, where the PONDR® VLXT score is the average disorder score of a query protein, and PONDR® VLXT (%) is a percent of the predicted disordered residues with the disorder scores above 0.5. Color blocks indicate regions in which proteins are mostly ordered (blue and light blue), moderately disordered (pink and light pink), or mostly disordered (red). If the two parameters agree, the corresponding part of background is dark (blue, pink, or red), whereas light blue and light pink reflect areas in which only one of these criteria applies. **B**. CH-CDF plot combining outputs of the charge-hydropathy (CH) and cumulative distribution function (CDF) analyses. The Y-coordinate is calculated as the distance of the corresponding protein from the boundary in the CH plot. The X-coordinate is calculated as the average distance of the corresponding protein’s CDF curve from the CDF boundary. The quadrant that the protein is located determines its disorder-based classification: Q1, protein predicted to be ordered by both tools (i.e., mostly ordered proteins); Q2, protein predicted to be ordered by CH-plot and disordered by CDF (i.e., native molten globules); Q3, protein predicted to be disordered by both tools (i.e., mostly disordered proteins); Q4, protein predicted to be disordered by CH-plot and ordered by CDF.

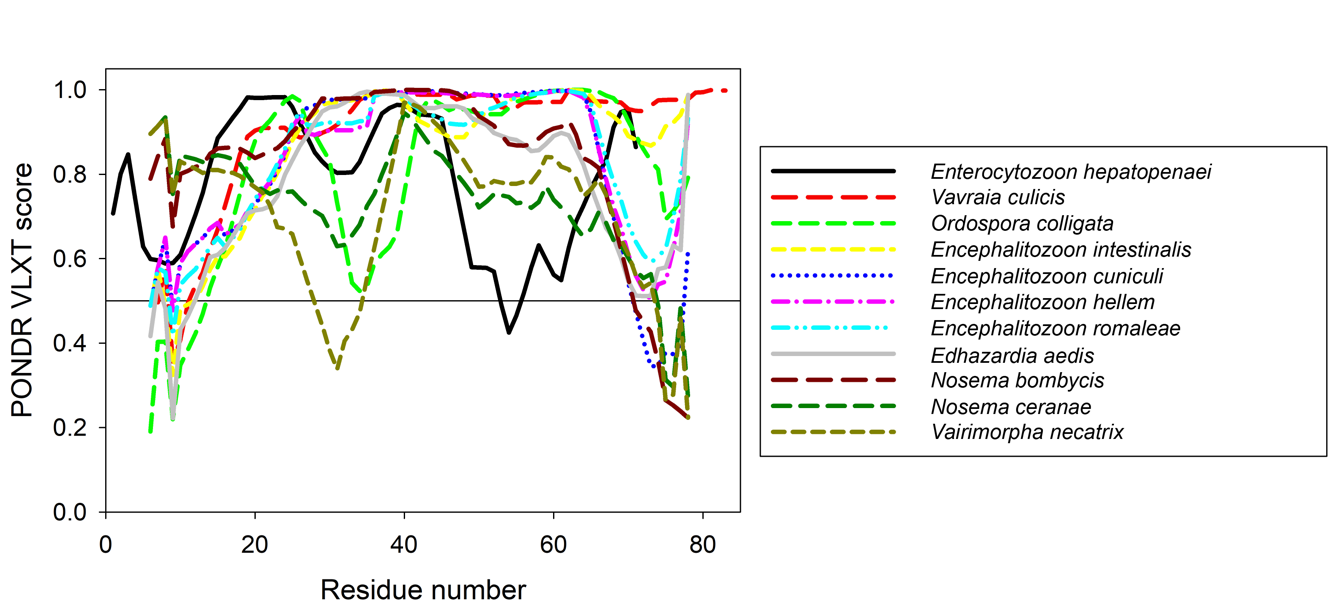

**Figure S3.** **Aligned intrinsic disorder profiles generated for the ribosomal proteins from the msL1/msL2 family by PONDR® VLXT predictor.** This plot compares homologous proteins the msL1/msL2 family in their intrinsic predisposition to remain unfolded (do not form a hydrophobic globule). In this plot, residues with the PONDR® VLXT scores exceeding 0.5 are expected to be disordered.
